# Ciliary propulsion and metachronal coordination in reef coral larvae

**DOI:** 10.1101/2022.09.19.508546

**Authors:** Rebecca N. Poon, Timothy A. Westwood, Hannah Laeverenz-Schlogelhofer, Emelie Brodrick, Jamie Craggs, Eric E. Keaveny, Gáspár Jékely, Kirsty Y. Wan

## Abstract

Larval dispersal is critical to the survival of coral reefs. As the only motile stage of the reproductive cycle, coral larvae choose a suitable location to settle and mature into adult corals. Here, we present the first detailed study of ciliary propulsion in the common stony reef coral *Acropora millepora*. Using high-speed, high-resolution imaging, particle image velocimetry, and electron microscopy, we reveal the arrangement of the densely packed cilia over the larval body surface, and their organisation into diaplectic (transversely propagating) metachronal waves. We resolve the individual-cilium’s beat dynamics and compare the resultant flows with a computational model of a ciliary array, and show that this form of ciliary metachronism leads to near-maximal pumping efficiency.

---

Ciliary motion is a distinctive and widespread method of propulsion across many aquatic species [1, 2]. Many planktonic organisms have cilia either covering their whole body, or localised into rings or lobed structures called ciliary bands [3, 4]. Unlike in cells with only a few cilia, such as sperm, or the biflagellate alga *Chlamydomonas* [5], arrays of beating cilia often exhibit sustained, long-range coordination of their beat phase, known as a metachronal wave (MCW), generating fluid flows whose global properties depend on the coordination patterns [6]. Thus, by organising ciliary activity, these tiny organisms can feed, swim, avoid predators and regulate their position in the water column [7, 8].

Cilia are present in the larvae of many marine invertebrates. The larval stage is that unique and critical phase of an organism’s life cycle that precedes or disguises its ultimate, mature form - hence the etymology of the term ‘larva’ from the Latin for ‘*a ghost, demon or mask* ‘ [9]. The common reef-building coral *Acropora millepora*, which is a broadcast spawning coral, produces motile larvae [10] following a short period of pelagic development. These settle and mature into adult colonies (Fig. 1), within *∼*50 days [11]. A suitable choice of location is vital for post-settlement survival [12]. Larval swimming and settlement depends on a variety of external cues such as light and chemicals [13–15], although this behaviour is little-understood from either a biophysical or a signalling perspective. Swimming is achieved by the beating of multiple cilia, densely distributed over the entire body surface. Although the primary role of these cilia is motility, they have also been shown to be useful in protecting larvae from suspended sediments [16].

**FIG. 1.**
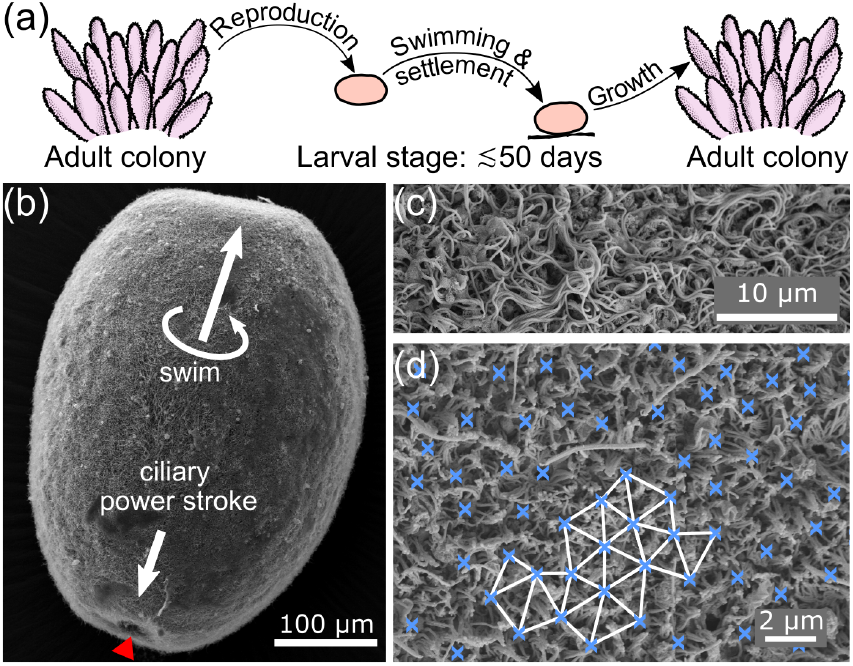
(a) Key stages of the coral reproductive cycle: an adult colony releases egg/sperm bundles, which combine to produce motile larvae. These eventually settle and metamorphose into an adult polyp, beginning a new colony. Scanning electron micrographs of (b) a whole *A. millepora* larva (oral pore marked by a red arrowhead); (c) the densely-ciliated surface; (d) a quasi-hexagonal arrangement of the cilia basal bodies in a deciliated preparation. Cilia bases (blue ‘x’) are surrounded by rings of microvilli.

Despite the obvious significance of coral larval motility for their survival to maturity, there has been no prior quantification of motility parameters or cilia behaviour. In this work, we present the first complete biophysical characterisation of ciliary motility and propulsion in a coral larva. We perform experimental measurements across scales, from flows induced by a single larva, down to the dynamics of individual cilia. Combining high-speed imaging and micromanipulation, we also resolve the beating waveforms of cilia, and show that surface cilia exhibit diaplectic metachronal coordination. Finally, we compare these data with a computational model of a densely ciliated array, to rationalise the likely functional significance of this ciliary coordination strategy for motility and flow pumping efficacy.

## Methods

*A. millepora* larvae were cultured *ex situ* [17, 18]. Annual spawning occurred during the winter of 2020/21 and live embryos (3 days post fertilisation) were transported directly from Horniman Museum and Gardens (London, UK) to an animal facility (University of Exeter, UK). Embryos were housed in artificial seawater containers inside incubators set to 27 °C. Larvae were fixed for scanning electron microscopy (SEM) in 2% gluteraldehyde at 4 dpf, dehydrated in ethanol and air dried in HMDS. After mounting on an aluminium sample pin with carbon conductive tab, they were sputter coated with 10 nm gold/palladium. Images of the larvae and their ciliated ectodermal surface were taken with a field scanning electron microscope (Zeiss GeminiSEM 500, Carl Zeiss Corp.) using a 5 kV beam. For flow field and particle image velocimetry (PIV) measurements, the fluid was seeded with 1 μm Fluoresbrite^®^ multifluorescent microspheres at a volume fraction of ≈ 10^−5^ v/v. Videos were post-processed using PIVlab [19]. Videos of individual free-swimming larvae in 500-600 μl droplets were captured at 80.5 fps using a Photometrics PRIME 95B camera mounted on a Nikon Ti2-U microscope, imaged with a 4X objective (CFI Plan Fluor) in autofluorescence (CoolLED pE300-ultra SB, 554/23 nm). Single-larva micromanipulation experiments were conducted using TW-1200 borosilicate glass micropipettes pulled using a P-1000 Puller (Sutter Instruments). Pipettes were scored and broken off to give an OD of (100 ± 10) μm, and fire polished to give a rounded tip with ID (35 ± 10) μm. High-speed images (100-1000 fps) were captured using a Phantom V1212 camera mounted on a Leica DMi8 inverted microscope, in brightfield or DIC.

## Experimental results

The larvae show a variety of body shapes, with individuals able to perform dramatic shape changes [20]. The basic shape is a prolate spheroid, with a slight narrowing and flattening at the oral pole. Larvae swim with the aboral pole facing forwards (Fig. 1b). Typical larval dimensions are length *∼* (785± 119) μm, width*∼* (404± 29) μm, with aspect ratios *α∼* 1.96± 0.37. Our SEM images show monociliated ectodermal cells, with each cilium surrounded by a ring of microvilli. The basal bodies show quasi-hexagonal order with lattice constant *a* =(1.8 ± 0.4) μm (Fig. 1c,d), so one larva has 𝒪(10^5^) cilia. The length of the motile, body cilia is *L* = (18 ± 2) μm. Since *a/L* = 0.1, comparable to *Paramecium*, where *a/L* = 0.15 [21], our system is a stereotypical ‘densely ciliated microswimmer’.

Videos of free-swimming larvae were sampled manually to find straight-swimming segments. The lab frame flows were measured for each video frame using PIV. The measured flow fields were shifted and rotated to place the larvae at the centre, and swimming towards the right hand side, of the frame, then averaged over at least one body rotation period to find the azimuthally-averaged flow field (see Video 1 and [20] for further examples). The larvae swim with a forward speed of (0.86 ± 0.06) mm s^−1^ and rotate about the body axis with a frequency of (0.23 ± 0.09) Hz. A full characterisation of the free-swimming behaviour will appear elsewhere [22]. Here we concentrate on details of the local, time-averaged flow fields around a freely swimming larva (Fig. 2a).

**FIG. 2.**
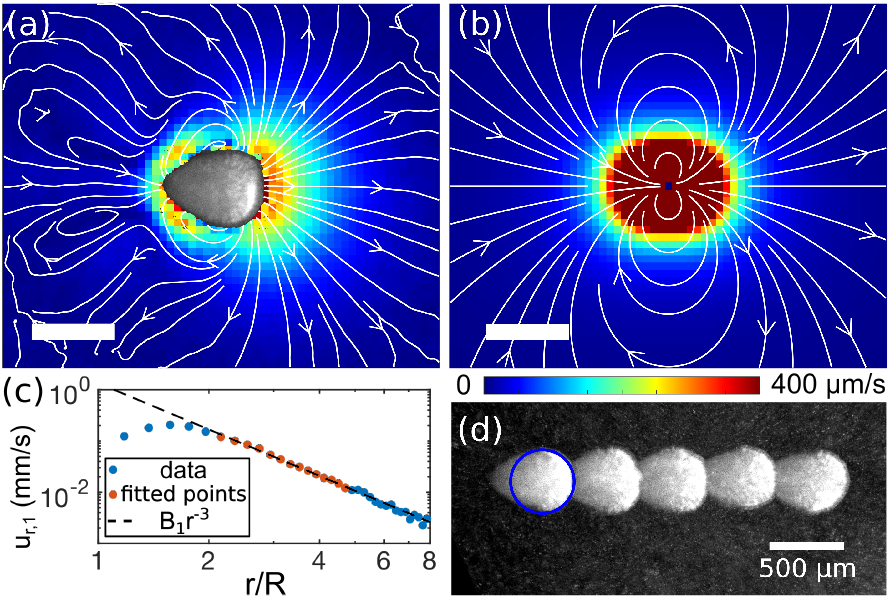
The flow field and streamlines around a free-swimming larva (a), compared with the fitted source dipole moment of the same flows (b). [The same color map was used in both cases. Scale bars: 500 μm.] (c) The first radial Legendre coefficient for our data. Orange points were used to fit the strength *B*_1_ of the source dipole moment, which decays as *r*^−3^. Radial distance *r* is normalised by the effective squirmer radius *R* for plotting clarity (see text). (d) Silhouettes of the free swimming larva at 0.5 s intervals. The blue circle shows the effective squirmer radius *R* =220 μm.

The coral larvae have a maximum forward speed of *U ∼* 1 mm s^−1^, and length 𝓁 ∼ 1 mm, so the Reynolds number for the larval motion is *Re* = *ρU*𝓁*/η* ≲ 1 (fluid density *ρ* and viscosity *η*). For the cilia (𝓁 ∼ 20 μm and maximum tip speed *U* ∼ 3 mm s^−1^), *Re* ∼ 0.6. Our hydrodynamics can thus be well described by the Stokes equations for incompressible fluid flow −∇*p* + *η∇* ^2^**u** = 0 and*∇ ·* **u** = 0, for fluid velocity **u** and pressure *p*.

Since the larvae reside at low *Re* and have high cilia density, we can use the ‘squirmer’ model [23, 24]. Squirmer motion is modelled by specifying a velocity at some continuous surface associated with the microswimmer, and applying a no-slip boundary condition. For a densely ciliated organism, this surface is the envelope covering the cilia tips, and the prescribed velocity represents the ciliary beat dynamics. In spherical coordinates with the origin at the centre of the larval body and *z* pointing in the swimming direction, the radial and tangential components *u*_*r*_ and *u*_*ϕ*_ of the fluid velocity of an azimuthally symmetric case are:

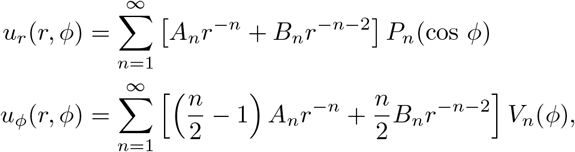

where *ϕ* is the polar angle, *P*_*n*_ are the Legendre polynomials, and

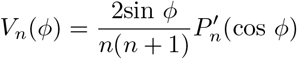

where ′ denotes the derivative with respect to cos *ϕ. A*_*n*_ and *B*_*n*_ can be calculated for a specified swimmer by matching the boundary conditions at the surface.

We estimate the leading order terms present in our measured flow fields by a numerical fitting procedure. Following [25], we calculated the Legendre coefficients of the radial flows [20], 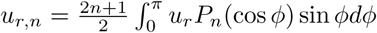. For *u*_*r*_ as given above, *u*_*r,n*_ = *A*_*n*_*r*_*−n*_ +*B*_*n*_*r*^*−n−*2^ (by completeness of the Legendre polynomials). For the larva in Fig. 2a, *u*_*r*,1_(*r*) is plotted in Fig. 2c. We observe a clear *r*^−3^ decay. The *r*^−1^ term is absent, so *A*_1_ = 0, consistent with a force-free swimmer. The source dipole moment has strength *B*_1_ = 1.5*×* 10^10^ μm^4^s^−1^. The flow field for a source dipole of this fitted strength (Fig. 2b) agrees reasonably well with the measured flows. Furthermore, a spherical squirmer with radius *R* and source dipole moment *B*_1_ has a swimming speed of *v*_0_ = 2*B*_1_*/*3*R*^3^ [25].Here, substituting our measured *v*_0_ = 890 μms^−1^ and fitted *B*_1_ = 1.5 *×*10^10^ μm^4^s^−1^ gives *R* = 220 μm. Although the larva is teardrop shaped, the fitted circle shows a good match to the size of the ‘head’ (Fig. 2d). Thus, to a first approximation the larva swims as a spherical squirmer with an effective radius of 220 μm.

To resolve the dynamics of the ciliated surface in detail, we tethered larvae to micropipettes in a variety of orientations. To specify these orientations, we now refer to a cylindrical coordinate system with **e**_*z*_ pointing in the larval swimming direction, and the azimuthal angle *θ* defined in a right-handed sense. First, we tethered the larvae ‘*sideways*’, with **e**_*z*_ in the image plane, Fig. 3a. The time-averaged flow field around a ‘*sideways*’ larva is shown in Fig. 3b. The maximum near-field tangential flow speed is similar to the maximum tangential flows measured in the comoving frame of the free-swimming larva, indicating that the native ciliary behaviour was not adversely affected by the pipette. The presence of vortices in the flow is likely due to confinement in the direction normal to the image plane [26].

**FIG. 3.**
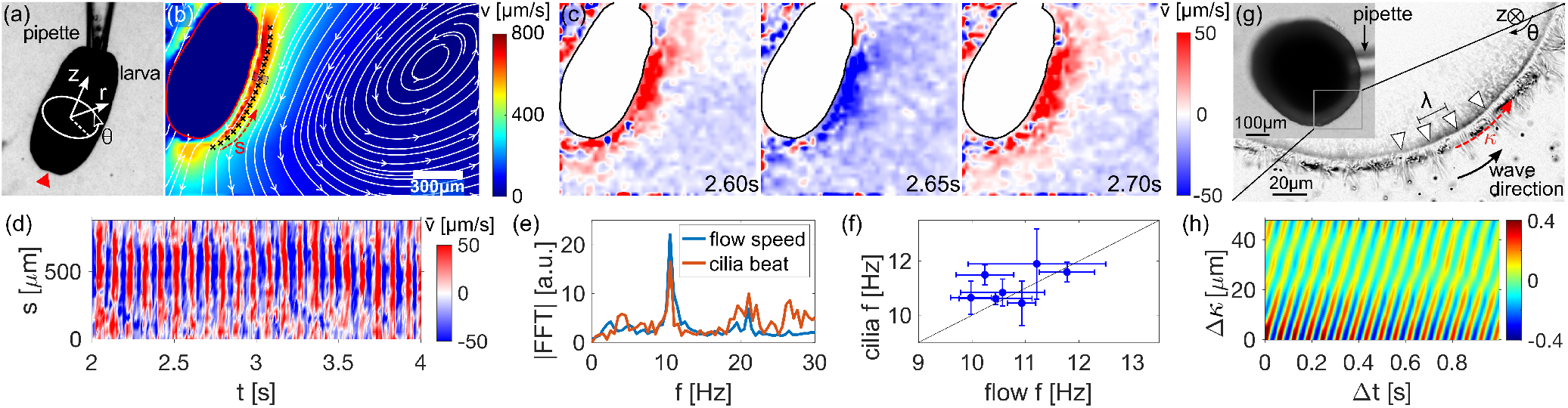
PIV performed on single coral larvae tethered by gentle micropipette suction at the aboral end (a). An example time-averaged flow field (b). (Streamlines shown in white. Colorbar indicates flow speed, and scale bar 300 μm.) (c) Three snapshots of the mean-subtracted flow speed 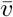 spaced at 50 ms. (d) Space-time kymograph of the near-field flow 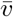 along the side of the larval body. Fourier transforms of the flow speed and the cilia beat peak at the same frequency for the pictured larva (e), and for the population (N=7) (f). (g) observation of a larva tethered in ‘top down’ configuration (**e**_*z*_ into the page) reveals metachronal wave crests. (h) Autocorrelation of cilia intensity along the ciliary band.

Using time-resolved flow fields as a proxy for the ciliary beat phase, we compute fluctuations around the mean flow speed, 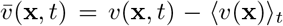. In the sideways orientation, 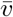 shows a periodic ‘blinking’ behaviour (Fig. 3c). The near field flow speed is plotted as a function of both time and arclength distance along the larval body in 3d. The ‘blinking’ is visible as vertical lines on the kymograph, indicating that the cilia beat *in-phase* along the aboral-oral axis. This is in contrast to PIV data from the model squirmer *V. carteri*, a spherical alga that shows alternating patches of high and low flow speed along the beat direction due to a MCW that propagates *in* the beat direction [27]. To confirm that the flow periodicity is related to ciliary beating, we compare the Fourier transform of the flow speed with of that of the cilia beat for the same video (from pixel intensity fluctuations at the surface): both show peaks at the same frequency of 10.5 Hz (Fig. 3e). Cilia beating matches the flow oscillation frequencies in 7 different larvae (Fig. 3f), for videos lasting ≳50 beat periods.

This implies a diaplectic MCW with vertical wave crests, propagating in either the +**e**_*θ*_ (*dexioplectic*, 90° anti-clockwise from the power stroke) or − **e**_*θ*_ direction (*laeoplectic*, 90° clockwise from the power stroke) [6]. To confirm the existence and direction of such a wave, we imaged the larvae in a ‘*top-down*’ state (Fig. 3g, Video 2). In this view, ciliary metachrony is clearly visible, and the wave propagates in the − **e**_*θ*_ direction. The 2D (time-space) autocorrelation of image intensity along the ciliary array as a function of time is used to estimate MCW parameters [28] (Fig. 3h). Wavecrests appear as diagonal lines, whose spacing in *κ* and *t* give the wavelength and period respectively (Fig. 3h). The wave was laeoplectic in all specimens where the wave direction could be deduced (*N* = 6). The cilia beat with frequency (11.1 ± 0.6) Hz, and the MC wavelength is (19 ± 4) μm.

## Computational results

Thus, the cilia are coordinated metachronally along the surface of the coral larva body in the direction shown in Fig. 4a. To reconcile the measured flow features with this putative ciliary coordination pattern, we developed a numerical simulation. Since the cilia are very short compared to the size of the larva (20–30 times), we approximate the curved ectoderm locally as a planar no-slip surface. A dense array of 1033 cilia of length *L* were arranged in a quasihexagonal lattice with spacing Δ_*θ*_ = 0.1*L* in the direction orthogonal to the beat plane, and Δ_*z*_ = 1.12*L* in the beat direction (the minimum allowable separation such that neighbouring cilia do not intersect). For each model cilium, its dynamics were prescribed according to high-speed video data (Video 3), and beating was assumed to be planar. Manually traced ciliary waveforms were simplified by fitting to a low-order polynomial in space and a four-term Fourier series in time to yield the trace shown in Fig. 4a, inset. Each cilium is discretized into *M* = 20 segments. For Stokes’ flow, the cilium segment velocities **V** and forces **F** are linearly related via the dense mobility matrix ℳ. The resulting system is solved numerically using a custom implementation of the Generalised Minimum Residual method [20, 29].

**FIG. 4.**
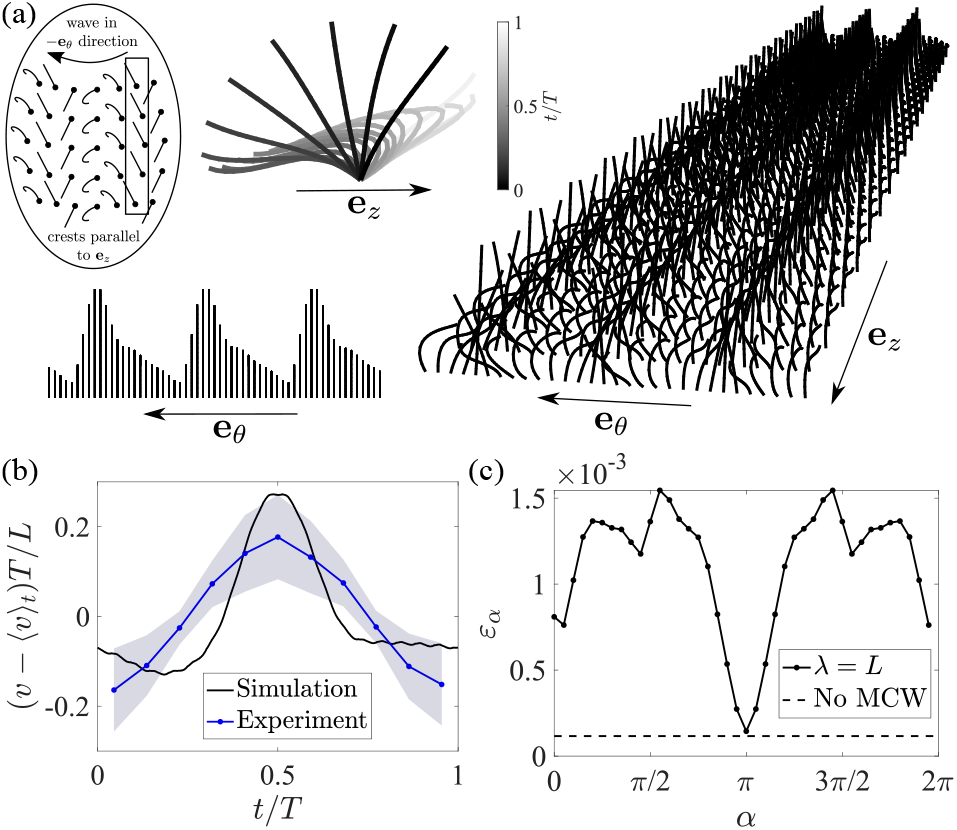
(a) Simulation snapshot of an array of 1033 model cilia, with ‘top-down’ view of the array and the model cilia waveforms over one cycle. Inset - schematic of the laeoplectic MCW on the larva (cilia not to scale). (b) Phase-dependent, mean-subtracted flow speed sampled in a box above the array in the simulations 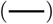 compared with experiments 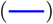 (N=7, average over multiple beat cycles. Shaded region: standard deviation). (c) The pumping efficiency of an array of model cilia for different MCW angles *α*.

At a height of *∼*5*L/*3 above the no-slip boundary, and midway along the larval body, we evaluated 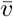 induced by the computational array when the cilia are coordinated in a laeoplectic wave, and compared this to the PIV sampled flow data (Fig. 4b). The simulations reproduced the periodic ‘blinking’ of the flows observed in Fig. 3b, where *T* is the beat period of the cilia. The quantitative agreement between the phase-dependent oscillatory flow profiles can be improved if technical limitations can be improved (e.g. the ability of PIV to resolve nearfield flows, inference of 3D motion from 2D microscopy), and/or if more realistic simulation parameters are chosen (even denser array, non-planar beat pattern).

Finally, we explore the functional consequences of different MCW directions for our computational ciliary array. To allow MCW waves of arbitary direction to propagate, Δ_*z*_ was increased to 1.76*L*. For different angles *α* = [0, 2*π*) between the MCW and the power stroke direction (*α* ={0, *π/*2, *π*, 3*π/*2}: {sym, dexio, anti, laeo}-plectic), we evaluated the dimensionless pumping efficiency 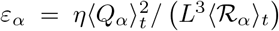 (Fig. 4c), where ⟨ *·*⟩ _*t*_ denotes a time-average, *Q*_*α*_ is the volume flow rate per cilium above the no-slip surface and ℛ_*α*_ = **V**^T^**F***/*1033 is the power exerted on the fluid per cilium [20]. The results reveal that the a metachronous state is always more efficient than perfectly synchronous beating, and diaplectic waves (travelling either to the left or right of the beat) are close to the global maximum in pumping efficiency.

## Concluding remarks

Reef-building corals, including *Acropora* species, have huge ecological importance yet have proved impossible to culture under laboratory conditions, until recently [18]. Here, we have presented the first biophysical study of cilia motility and coordination in a coral larva. Our multiscale approach covers individual cilium dynamics, their collective coordination, and cilia-induced flows around whole larvae. We found that the density of cilia in the ectoderm is high, allowing comparisons with a squirmer model [23, 24]. For larvae with aspect ratio close to unity, the free-swimming flows are well-described by a spherical squirmer whose (effective) radius (obtained by our flow-field matching procedure) is comparable to the size of the larval ‘head’. This is in spite of the deviation of larval morphology from perfect spheres, and further breaking of azimuthal symmetry by both cilia orientation and coordination.

The larvae rotate slowly about the aboral-oral axis whilst swimming. This may arise from slightly nonplanar ciliary beating, as seen in *Chlamydomonas* [5]. Here, we neglected azimuthal asymmetries by averaging free-swimming flows over multiple rotation periods, justified since the period of cilia beating (*∼*0.1 s) is much smaller than that of body rotation (*∼*2 s). We can naturally extend the spherical squirmer model to include azimuthal asymmetry [30], or more realistic spheroidal or rod-shaped larval morphologies [25], thereby coupling single-cilium dynamics with whole-larva propulsion [22].

We also make the first observation of MCWs in a coral larva. By imaging body-fixed specimens in multiple viewing angles, we deduced that the wave is diaplectic (particularly, laeoplectic). In most other organisms where MCWs have been unambiguously characterised they have been either symplectic [6, 31], or antiplectic [32, 33]. Generally, some form of metachronism in ensembles of cilia is to be expected due to hydrodynamic interactions [34–36], but the wave propagation direction remains highly system-dependent and ultimately unexplained in living organisms. Insights from idealized scenarios, both simulated and artificial, emphasise the importance of the cilia beat pattern, organisation, and boundary conditions [29, 37–40]. Ciliary coordination could even be influenced by interactions with mucous, as shown in human bronchial epithelia [41]. The larvae of the coral *Caryophyllia smithii* trail mucous strings from the oral pore to facilitate feeding [42]. Although it is not known whether *A. millepora* larvae are similar feeders, we did observe ‘mucous trails’ being dragged behind the swimming larvae. Mucous secretion [16] hinders our ability to perform near-field PIV (could be the cause of the two ‘lobes’ in Fig. 2a). Thus, to answer the question of how diaplectic waves actually *emerge* in the coral ectoderm, more experiments and modelling are needed.

Diaplectic metachronism has been described anecdotally in diverse invertebrate larvae [6, 43]. The sense of MCWs, if they exist, should be inferred carefully in other cnidarians. Here we used a realistic simulation of a densely-ciliated array and beat patterns reconstructed from data to show that diaplectically-directed MCWs are close to optimal for pumping efficiency. In the stroke direction, steric interactions promote in-phase synchronization since interciliary spacing is so small (0.1*L*); in the transverse direction however, phase shifts help prevent collisions between closely-packed beating cilia [44]. From protists to humans, the organisation and patterning of cilia undergoes precise developmental control [28, 45], so improved pumping efficiency due to diaplectic metachronism could be a key evolutionary driver for ciliary placement and basal-body orientation.

This work was funded by the Human Frontier Science Programme grant RGP0033/2020 (G.J.), an EPSRC Doctoral Prize Fellowship [EP/T51780X/1] (T.A.W), and the European Research Council (ERC) under the European Union’s Horizon 2020 research and innovation programme grant 853560 EvoMotion (K.Y.W). To induce *ex situ* coral spawning, reagents and aquarium filtration equipment were provided by Triton Applied Reef Bioscences and EcoTech Marine (J.C.). We also acknowledge support from the Wolfson Bioimaging facility at the University of Bristol.

