## Supplemental Material for "Ciliary propulsion and metachronal coordination in reef coral larvae"

[1]*Living Systems Institute, University of Exeter, Exeter EX4 4QD, United Kingdom*
[2]*Department of Mathematics, Imperial College London, London SW7 2AZ, United Kingdom*
[3]*Horniman Museum and Gardens, London, SE23 3PQ, United Kingdom*
(Dated: September 19, 2022)

### S1. DIVERSITY OF OBSERVED LARVAL SHAPES

The larvae possess musculature allowing them to change shape over time, similar to the larvae of other cnidarians [1]. Additionally, there is a high degree of variation between the shape of different individuals. Figure 1 shows a gallery of observed larval shapes as imaged in our micropipette experiments.

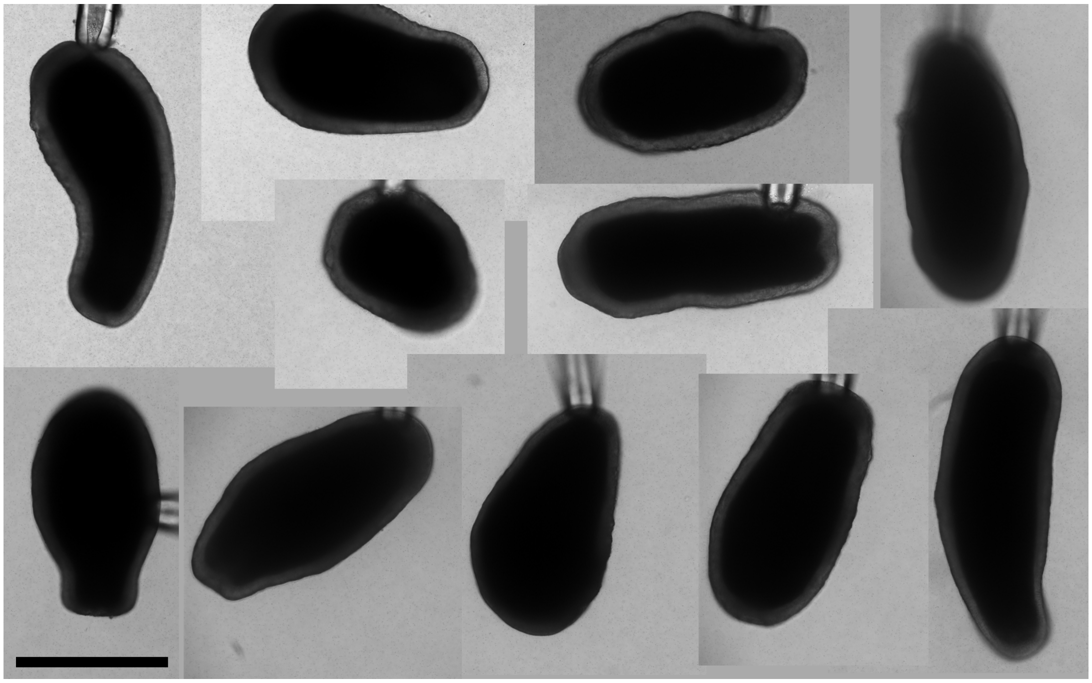

FIG. 1. Observed shapes of 9 different larvae (a-i). Views at two different times are shown for larvae (a) and (c), showing their ability to change shape change over time. Scale bar: 500 µm.

### S2. FITTING LEGENDRE COEFFICIENTS

The Legendre coefficients for fitting the source dipole from measured flow fields are calculated by evaluating the integral

$$u_{r,n}(R) = \frac{2n+1}{2} \int_0^{\pi} u_r(R,\phi) P_n(\cos\phi) \sin\phi d\phi. \tag{1}$$

over one hemisphere (over $0 < \phi < \pi$), at values $r = R$ between 0 and the edge of the video frames. $u_r(R, \phi)$ is the radial component of the flow field at the point $(R, \phi)$, and $P_n$ is the $n$th Legendre polynomial. We evaluate the integral numerically from our data. Figure 2 shows a schematic of the points used to evaluate the sum.

The centre of the larva is used for the origin of the radial coordinate system. To define the centre of the larva, we threshold the images to locate the body, and use the centre of the bounding rectangle of the thresholded particle.

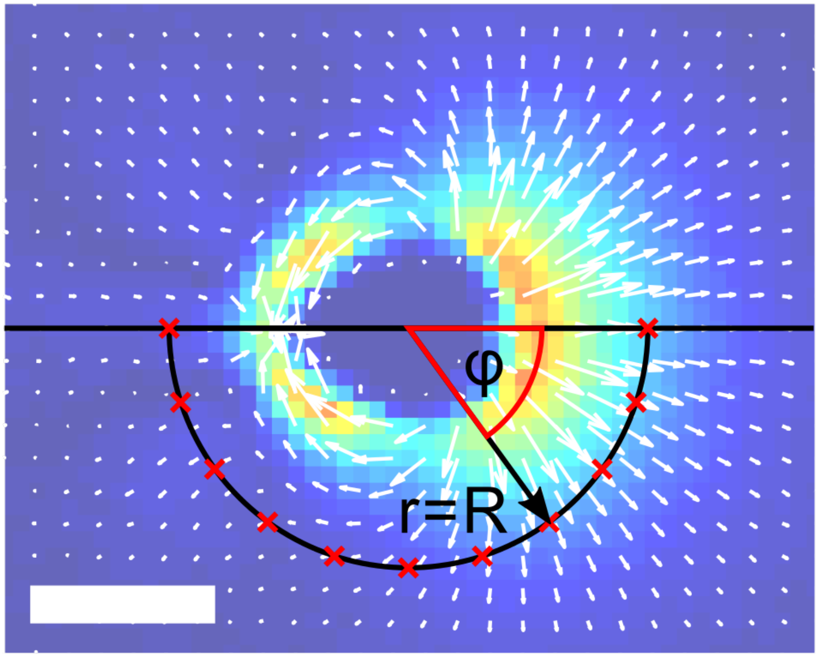


FIG. 2. Schematic for Legendre coefficient fitting. The integrand is evaluated at some $r = R$, at points (red crosses) that are evenly spaced in $\phi$, then summed over $\phi$ to find $u_{r,n}(R)$. Scalebar: 500 µm.

### S3. FURTHER EXAMPLES OF SINGLE-LARVA FLOW FIELDS

The analysis used for the larva shown in Figure 2 was performed on 3 other larvae. Fig. 3 shows the experimentally measured flow fields, overlaid with the fitted squirmer sizes, which were calculated using the same method as given in the main text. The larvae were fitted to spherocylinders with aspect ratios of 1, 1.5 and 2, using the correction factor for aspect ratios $> 1$ given in [2]. The shapes and sizes of the fitted squirmers show varying degrees of similarity to those of the larvae. Thus whilst the squirmer description is valid as a first approximation, it is not always a complete description of the swimming.

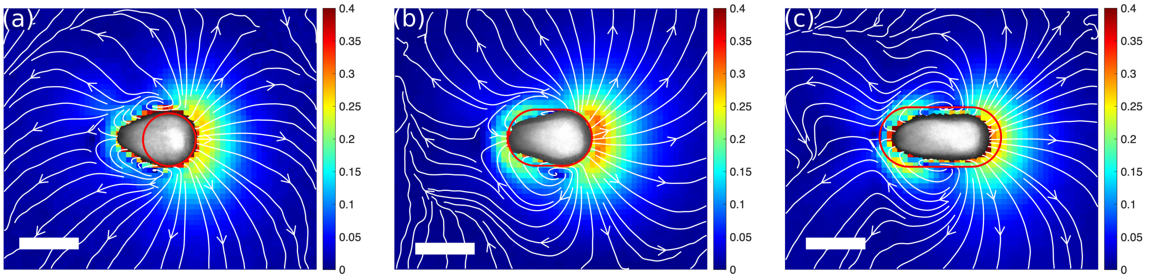


FIG. 3. Experimental free swimming flow fields for three more larvae, with fitted squirmer sizes overlaid. (a) was fitted with a sphere, and (c,d) with spherocylinders of aspect ratio 1.5 and 2 respectively. Scalebars: 500 µm.

### S4. ESTIMATING FREQUENCIES OF OSCILLATORY FLOWS AND CILIARY BEATING

In the ‘sideways’-view, oscillatory flow frequency was calculated from kymographs as shown in Figure 3b. The Fourier transform of $v(t) - \langle v \rangle$ was calculated for each box (i.e. each row of the kymograph), then averaged. For each video, the peak location and the full width at half maximum was then calculated to give the mean frequency and the error bar respectively, as shown in Figure 3f. For the individual presented in Figure 3(a-f), the cilia beat signal was calculated from the same 10x, sideways oriented video that was used to calculate the flow data (as described in the main text). For one additional larva, a top-down video was not available, so a 40x magnification, sideways video was used. For the five remaining larvae, we used top down videos of the wave at 40x objective magnification for the CBF estimation. Cilia beat frequency was calculated from high-speed videos of the ciliated ectoderm, by measuring the fluctuation in the pixel standard deviation (larvae 1 and 2) or mean intensity (larvae 3-7) within small ROIs located in the ciliary band [3], then taking a Fourier transform of the fluctuating signal. We observed a possible dependence of the cilia beat frequency on light levels, but further experiments would be necessary to verify this.

### S5. FOURIER-FITTING CILIA BEAT WAVEFORMS

Stereotypical cilia beat patterns were found by tracing over videos of beating cilia by hand (Video 2) using the ‘freehand line’ function in ImageJ, and exporting the lines as coordinates. This was done for three different cilia, with traces spaced at 4 ms intervals. Figures 4a-c show the three independent traces. In all cases, it is clear that the cilia are extended perpendicularly relative to the body surface during the power stroke, and display a high-degree of curvature during the recovery stroke when they move almost parallel to the body surface.

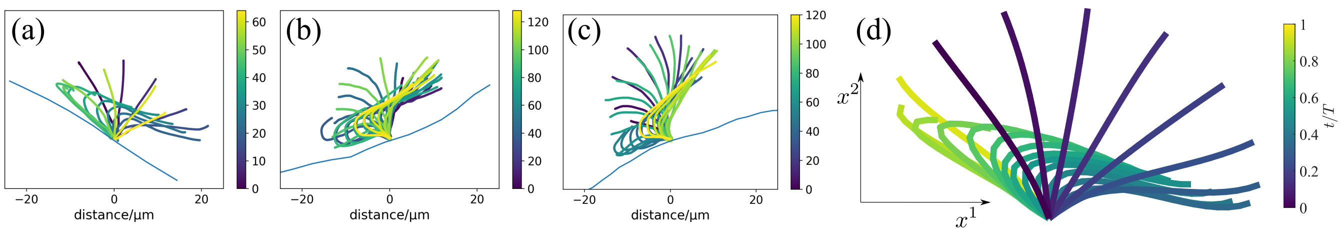


FIG. 4. (a-c) Beat traces for three cilia. The traces are colored according to time, and the colorbar shows time in ms. The traces are spaced at 4 ms intervals. (d) The simulation waveform fitted to the trace in (a) over the course of one period $T$.

For the cilia-array simulations, we generate analytic approximations to the measured beat patterns by performing a Fourier-series-least-squares fit to these traces, as described by Blake [4]. Specifically, to obtain the Fourier-fitted tracing in Figure 4d, we first identify 20 points equally spaced along the length of the trace at each instant in time. Note that because they are two-dimensional images of a three-dimensional motion, the arclength of the traces varies slightly over time and so we also scale each trace to have unit length at this stage. For each of these 20 points, we then fit a truncated Fourier series with $N$ modes to the point’s position over time. Finally, we fit cubic polynomials to the Fourier coefficients (in a least-squares sense) as they vary along the cilium length. The result is a collection of constant coefficients, $A^i_{nm}$ and $B^i_{nm}$, such that the $i$-th component, $i \in \{1, 2\}$, of the position of a point $s \in [0, 1]$ along a cilium of length $L$ at time $t$ is approximated as

$$x^i(s,t) = L \sum_{n=0}^{N-1} \sum_{m=1}^{3} \left( A^i_{nm} \cos(2n\pi t/T) + B^i_{nm} \sin(2n\pi t/T) \right) s^m. \tag{2}$$

For the traces in this work, we used $N = 4$ Fourier modes.

Note that because of the fitting procedure, the length of a model cilium described by eq. (2) is not constant over time (see fig. 5), much like the original experimental data. However, the coefficients, given in table I, are scaled such that the average length of the cilium over time is $L$. The resulting waveform as a result of applying this procedure to the experimental traces in fig. 4a can be seen in fig. 4d.

Code for decomposing experimentally-tracked ciliary waveforms into their respective Fourier modes can be found at `https://github.com/timwestwood/fit_cilium_beat_pattern_to_data`.

### S6: SIMULATION METHODS

To further investigate the implications of the metachronal coordination exhibited by the cilia, we performed numerical simulations of 3D model cilia whose shapes are described by eq. (2) (up to some cilium-specific phase shifts $2\pi t/T \mapsto 2\pi t/T + \varphi_0$, $\varphi_0 \in [0, 2\pi)$, which encode the coordinated motion). In all simulations discussed in this work, we arranged a total of 1033 model cilia on a no-slip planar boundary according to an irregular hexagonal tiling. Each

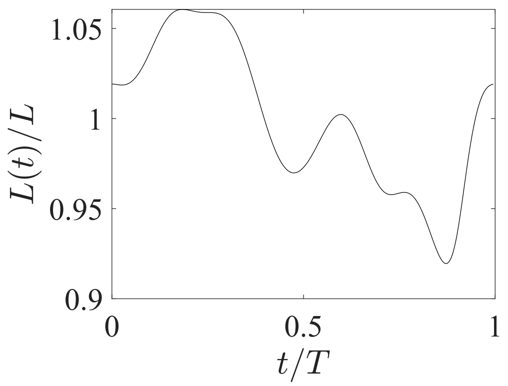


FIG. 5. The length of a model cilium described by eq. (2) over the course of one period $T$.

| $A^1_{mn}$ | | $n$=0 | 1 | 2 | 3 |
|---|---|---|---|---|---|
| $m$ | 1 | -0.2766 | 0.3091 | -0.2952 | -0.0987 |
| | 2 | 0.6369 | -1.9937 | 0.7189 | 0.3313 |
| | 3 | -0.0851 | 1.0918 | -0.6329 | -0.2578 |

| $A^2_{mn}$ | | $n$=0 | 1 | 2 | 3 |
|---|---|---|---|---|---|
| $m$ | 1 | 0.9034 | -0.2469 | 0.3076 | 0.0238 |
| | 2 | -0.6635 | 1.2159 | -0.6966 | -0.1356 |
| | 3 | 0.2016 | -0.5867 | 0.4555 | 0.0732 |

| $B^1_{mn}$ | | $n$=0 | 1 | 2 | 3 |
|---|---|---|---|---|---|
| $m$ | 1 | 0 | 0.9166 | -0.2946 | 0.0612 |
| | 2 | 0 | 0.0885 | 1.0580 | -0.2319 |
| | 3 | 0 | -0.4839 | -0.6336 | 0.1906 |

| $B^2_{mn}$ | | $n$=0 | 1 | 2 | 3 |
|---|---|---|---|---|---|
| $m$ | 1 | 0 | 0.1337 | 0.0815 | 0.0362 |
| | 2 | 0 | -0.6808 | 0.3473 | 0.0862 |
| | 3 | 0 | 0.6009 | -0.2744 | -0.0612 |

TABLE I. The coefficients required for the model cilium shape given in eq. (2).

cilium is prescribed to follow a planar beat pattern. The base-to-base separation orthogonal to the beat plane was set to $0.1L$ to match the values reported in experiments, but the spacing parallel to the cilia beating is chosen to prevent the model cilia from intersecting during the simulations.

In the simulation designed for direct comparison to the experimental flow field measurements (see fig. 4b in the main text), the prescribed laeoplectic metachronal wave guarantees that all model cilia are in-phase with their beat-wise neighbours, and we can use a separation of $1.12L$. However when we vary the angle, $\alpha$, between the effective stroke and the wave's propagation direction (for fig. 4c in the main text) we must allow for arbitrary phase differences between beat-wise neighbours and instead use a slightly larger separation of $1.76L$ to prevent steric interactions. See fig. 6 for an illustration of this geometry. Note also that to achieve an angle $\alpha$ between the stroke and wave directions, as well as a wavelength of $\lambda$, a model cilium whose base has position $X_\theta \mathbf{e}_\theta + X_z \mathbf{e}_z$ is given initial phase-shift $\varphi_0 = 2\pi(X_z \cos(\alpha) - X_\theta \sin(\alpha))/\lambda$. In all simulations presented in this work we use $\lambda = L$ to match the experimental data.

At each given time $t$, we consider $M = 20$ discrete points along the length of each model cilium. In order to accurately resolve the shape and motion of the cilium, we choose the $m$-th point, $1 \le m \le M$, to be at a fraction $l_m = (m-1)/(M-1)$ along the length of the model cilium. As a result of the fitting procedure, the parameter $s \in [0,1]$ in eq. (2) does not represent the length fraction along the cilium and we cannot simply use $s_m = l_m$ to define the location of the $m$-th point through eq. (2) (except for the basal ($m = 1$) and distal ($m = M$) points). Instead, the appropriate $s_m$ are solved for numerically for each shape the model cilium attains.

Having done so, we can define the velocity of each point as the time derivative of eq. (2) evaluated at the appropriate $s_m$; let the vector $\mathbf{V} \in \mathbb{R}^{3099M}$ denote the collection of the velocities of all $M$ points in all 1033 model cilia. As discussed in the main text, the hydrodynamics of the coral larvae can be described by the incompressible Stokes equations,

$$\begin{aligned} \mathbf{0} &= \eta \nabla^2 \mathbf{u} - \boldsymbol{\nabla} p, \\ 0 &= \boldsymbol{\nabla} \cdot \mathbf{u}, \end{aligned} \tag{3}$$

where $\mathbf{u}$ is the fluid velocity, $p$ is the pressure and $\eta$ is the fluid viscosity. This yields a linear relationship between

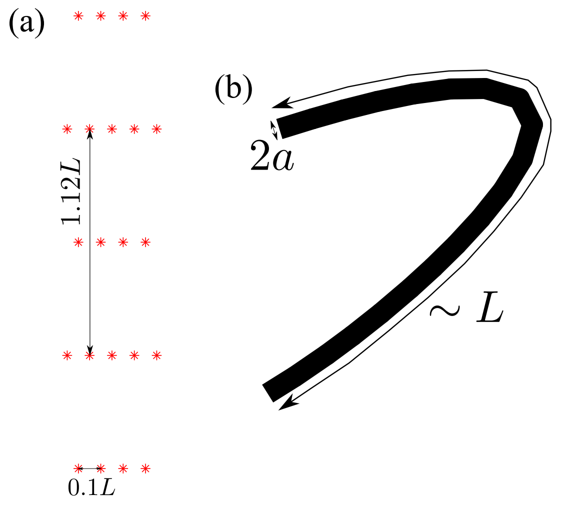


FIG. 6. (a) A schematic of the irregular hexagonal cilium placement used in the flow speed oscillation simulation (see fig. 4a-b in the main text). For the $\alpha$-sweep simulations discussed in fig. 4c of the main text, the geometry is the same with $1.12L$ replaced by $1.76L$. (b) The geometry of a model cilium; recall that the length scale $L$ is the average length of the model cilium over a single period.

these velocities, $\mathbf{V}$, and the collection of forces exerted by the model cilia on the fluid at these points, $\mathbf{F} \in \mathbb{R}^{3099M}$, which is described by the so-called mobility matrix $\mathcal{M} \in \mathbb{R}^{3099M \times 3099M}$; i.e.

$$\mathbf{V} = \mathcal{M}\mathbf{F}. \tag{4}$$

The mobility matrix describes the hydrodynamic interactions between all discrete points in all model cilia, and in this work it is approximated using the modifications to the Rotne–Prager–Yamakawa mobility for particles above an infinite no-slip boundary described by Swan and Brady [5]. Note that this mobility approximation requires that we assign a radius, $a$, to each discrete point. We nominally identify this radius with the radius of the cilium cross-section (see fig. 6), and in this work we use the aspect ratio $a/L = 0.02125$.

Thus at each instant in time we can solve the linear system eq. (4) for the forces $\mathbf{F}$. We do so iteratively using a custom C++ implementation of the GMRES method [6], with the multiplications by $\mathcal{M}$ accelerated on GPUs by custom CUDA kernels [7, 8]. Once we have the forces, we can evaluate other quantities of interest such as the rate of viscous dissipation (i.e. the power exerted on the fluid) per cilium, $\mathcal{R} = \mathbf{V}^\top \mathbf{F}/1033$.

For example, the flow velocity $\mathbf{u}$ at a point $\mathbf{x} = (x_1, x_2, x_3)^\top$ in the fluid can be evaluated by summing over all forces contained in $\mathbf{F}$ – $\mathbf{F}_k \in \mathbb{R}^3$, $1 \leq k \leq 1033M$ – the corresponding Green's function solution for the flow above a no-slip boundary, often called the Blakelet solution [9]:

$$\begin{aligned}\mathbf{u}(\mathbf{x}) = \frac{1}{8\pi\eta} \sum_{k=1}^{1033M} \Bigg\{ & \left( \frac{1}{\|\mathbf{r}_k\|} - \frac{1}{\|\mathbf{R}_k\|} \right) \mathcal{I} + \frac{\mathbf{r}_k \mathbf{r}_k}{\|\mathbf{r}_k\|^3} - \frac{\mathbf{R}_k \mathbf{R}_k}{\|\mathbf{R}_k\|^3} \\ & + \frac{2 y_k^{(3)}}{\|\mathbf{R}_k\|^5} \left( \mathbf{R}_k \mathbf{R}_k \cdot \begin{bmatrix} 3x_3 & 0 & 0 \\ 0 & 3x_3 & 0 \\ 0 & 0 & -3x_3 \end{bmatrix} + \|\mathbf{R}_k\|^2 \begin{bmatrix} -x_3 & 0 & r_k^{(1)} \\ 0 & -x_3 & r_k^{(2)} \\ r_k^{(1)} & r_k^{(2)} & x_3 \end{bmatrix} \right) \Bigg\} \cdot \mathbf{F}_k,\end{aligned} \tag{5}$$

where $\mathbf{y}_k = \left( y_k^{(1)}, y_k^{(2)}, y_k^{(3)} \right)^\top$ is the point at which the force $\mathbf{F}_k$ is applied to the fluid, $\mathbf{r}_k = \left( r_k^{(1)}, r_k^{(2)}, r_k^{(3)} \right)^\top = \mathbf{x} - \mathbf{y}_k$ is the displacement from this point and $\mathbf{R}_k = \left( r_k^{(1)}, r_k^{(2)}, r_k^{(3)} + 2y_k^{(3)} \right)^\top$ is the displacement from the image of this point across the no-slip boundary at $y^{(3)} = 0$. The flow speeds discussed in fig. 4b of the main text are then simply the magnitudes of these velocities (with the component out of the beating plane discarded), averaged over 5000 equally-spaced points in a rectangle of dimensions $2.5L$-by-$5L$ whose base is at a height $5L/3$ above the wall to match the experimental procedure.

We also evaluate the pumping efficiency, $\varepsilon$, of arrays of model cilia. We define the pumping efficiency very similarly to Guo *et al.* [10] as

$$\varepsilon = \frac{\eta \langle Q \rangle_t^2}{L^3 \langle \mathcal{R} \rangle_t}, \tag{6}$$

where $\mathcal{R}$ is the per-cilium rate of viscous dissipation defined above and

$$Q = \frac{1}{1033\pi\eta} \sum_{k=1}^{1033M} y_k^{(3)} \left( \mathbf{F}_k \cdot -\mathbf{e}_z \right) \tag{7}$$

is the per-cilium volume flow rate in the flow (i.e. $-\mathbf{e}_z$) direction through a half-plane above the no-slip boundary [11, 12].

### LIST OF SUPPLEMENTARY MOVIES

*Movie 1: video of a free-swimming coral larva* The video was taken in autofluorescence (554/23 nm wavelength), using a 4x magnification objective lens. The fluid is seeded with fluorescent beads to allow measurement of the flow. The larva swims in a straight path, turning about its own axis.

*Movie 2: metachronal wave from top-down view* Video taken in DIC microscopy with a 40x magnification objective lens, and processed by subtracting the background of each frame. The cilia beat towards the viewer, so the wave is laeoplectic. (Due to the fact that images are inverted in the microscope, the video has been flipped so that the orientation and handedness are consistent with an observer viewing the sample from below.)

*Movie 3: manual tracing of a single coral cilium* Video taken in DIC, with a 63x magnification objective lens. The position of one cilium was traced by hand in successive frames, at 250 fps.

*Movie 4: simulation of a metachronal wave* A simulation of a dense 2D array of cilia. The beat shape and frequency are prescribed, and the relative phases are of neigbouring cilia are prescribed to give a laeoplectic metachronal wave.

---
